# Finally, Bulk Typing of Bacterial Species down to Strain Level using ON-rep-seq

**DOI:** 10.1101/402156

**Authors:** Łukasz Krych, Josué L. Castro-Mejía, Daniel N. Moesby, Morten B. Mikkelsen, Morten A. Rasmussen, Maciej Sykulski, Dennis S. Nielsen

## Abstract

Despite the massive developments within culture-independent methods for detection and quantification of microorganisms during the last decade, culture-based methods remain a cornerstone in microbiology. We have developed a new method for bacterial DNA enrichment and tagmentation allowing fast (< 24h) and cost-effective species level identification and strain level differentiation using the MinION portable sequencing platform (ON-rep-seq). DNA library preparation takes less than 5h and ensures highly reproducible distribution of reads that can be used to generate strain level specific read length counts profiles (LCp). We have developed a pipeline that by correcting the random error of reads within peaks of LCp generates a set (∼10 contigs per sample; 300bp - 3Kb) of high quality (>99%) consensus reads. Whereas, the information from high quality reads is used to retrieve species level taxonomy, comparison of LCp allows for strain level differentiation. With benchmarked 288 isolates identified on a single flow cell and a theoretical throughput to evaluate over 1000 isolates, our method allows for detailed bacterial identification for less than 2$ per sample at very high speed.

## Introduction

Culture dependent methods remain indispensable in detailed identification of bacteria. Yet, successful typing of bacteria down to species/strain level remains not fully resolved ^1^. Several promising technologies and methodologies for solving the problem have been proposed but with a variable success. Generally, fast and cost-effective methods are not accurate enough, while those that are more accurate are also more laborious and/or expensive. Methods based on 16S rRNA gene sequencing are amongst the most universal, yet species level resolution cannot always be reached ^2^. More complex molecular tools that are able to reach strain level resolution such as PFGE, Rep-PCR, MLST or MALDI-TOF MS are hampered by one or several drawbacks that include low speed/throughput, limited databases, no taxonomic information, laborious procedure or high equipment cost ^3,4,5^.

The present gold standard for strain level bacterial identification is full genome sequencing. Optimally this approach combines information from high-throughput, short, good quality reads with lower throughput, poor quality but long reads ^6^. However, this approach is far from being cost effective, and the data analysis and interpretation is far from trivial ^7,8,9^.

The portable DNA sequencing platform MinION by Oxford Nanopore Technologies (ONT) offers an attractive tool with a potential to tackle the task of species/strain level identification ^10^. Unfortunately, ONT still deals with two critical problems: relatively high error rate at the base level and lower throughput compared to technologies offered by e.g. Illumina ^10^. We propose a DNA enrichment method that to a large extent have solved both these pitfalls by combining an optimized version of repetitive extragenic palindromic PCR (Rep-PCR) with a consecutive dual-stage Rep-PCR-2 step during which sample specific barcodes are incorporated.

Repetitive extragenic palindromic sequences in bacterial genomes were first described in the genomes of *E. coli* and *Salmonella* in 1984 by M.J. Stern ^11^. A decade later J. Versalovic used interspersed repetitive sequences as a binding site for primers developing Rep-PCR ^12^. Amplicons varying in length (from few dozens base pairs (bp) to few kilo base pairs (Kbp)) separated with electrophoresis create a genomic fingerprint that has been proven many times to have species and in some set-ups also strain level discriminative resolution of bacteria ^13^. Only five years later Rep-PCR was described as one of the most reproducible and commonly used method for species and strain level discernment ^14^, and numerous applications of the method have been reported in many fields including food processing, food safety, environmental microbiology, and medicine ^15,16,17,18,19^. Despite the immense progress in DNA sequencing technologies Rep-PCR is still a commonly used technique in many research groups mainly due to the low cost of the analysis and basic laboratory equipment needed ^20^. However, the low running costs comes with a price of highly laborious and time-consuming procedures involving 3-5h PCR, 3-5h electrophoresis, and complicated, tedious and potentially error prone fingerprint data analysis. Additionally, Rep-PCR only allows for bacterial discrimination but not direct identification ^21^.

We are presenting a new bacterial DNA enrichment method for Oxford Nanopore sequencing called ON-rep-seq. The method exploits an optimized version of Rep-PCR for reproducible amplification followed by a dual stage Rep-PCR-2 step allowing tagmentation of up to 96 samples in one reaction. Furthermore, we have developed a pipeline utilizing the information from the generated sequences at three levels: i) generation and comparison of isolate specific read length counts profiles (LCp) ii) detection of peaks in each LCp followed by within-peak correction of the random single base error iii) species-level taxonomy assignment using corrected consensus reads (Figure 1). The method has been tested on 38 different bacterial species and three strain level groups successfully identifying all bacteria down to the species level and discriminating strains with a sensitivity that is at least similar to a Whole Genome Sequencing (WGS) based approach.

**Figure 1.**
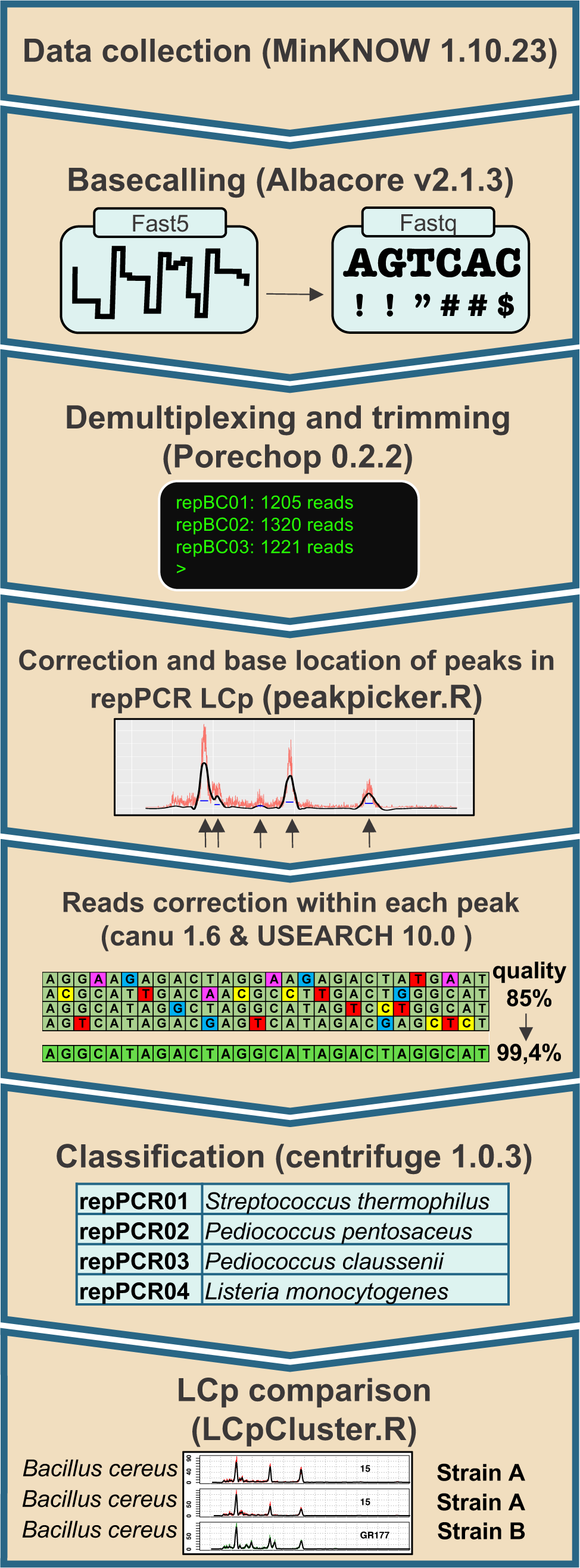
ON-rep-seq pipeline overview. The schema describing pipeline that allows processing of the raw Oxford Nanopore Technology based rep-PCR amplicon sequencing (ON-rep-seq) data. After initial besecalling, demultiplexing (separating according to barcodes) the fastq files are used to generate read length counts profiles (LCp) based on sequences length distribution. Subsequently, reads within each peak are clustered with USEARCH, corrected with Canu, followed by Centrifuge based taxonomy classification using improved quality reads. Finally, the traces can be compared to estimate strain level relatedness between pairs of LCp.

## Results

### Sequencing of the Rep-PCR enriched library with MinION generates highly reproducible LCp

Similar to Rep-PCR gel based fingerprints, sequenced Rep-PCR products can be transformed into read length counts profiles (LCp) being a function of reads length and abundance. The shape and position of peaks is highly reproducible in all technical replicates across first two sequencing runs (Figure 2, Supplementary Figures 1 and 2) indicating that the barcode sequences do not affect the shape or the position of the peaks during Rep-PCR-2. Yet, as explained below, we observed a minor run effect in the third consecutive run resulting in shifted distribution of short/long reads.

**Figure 2.**
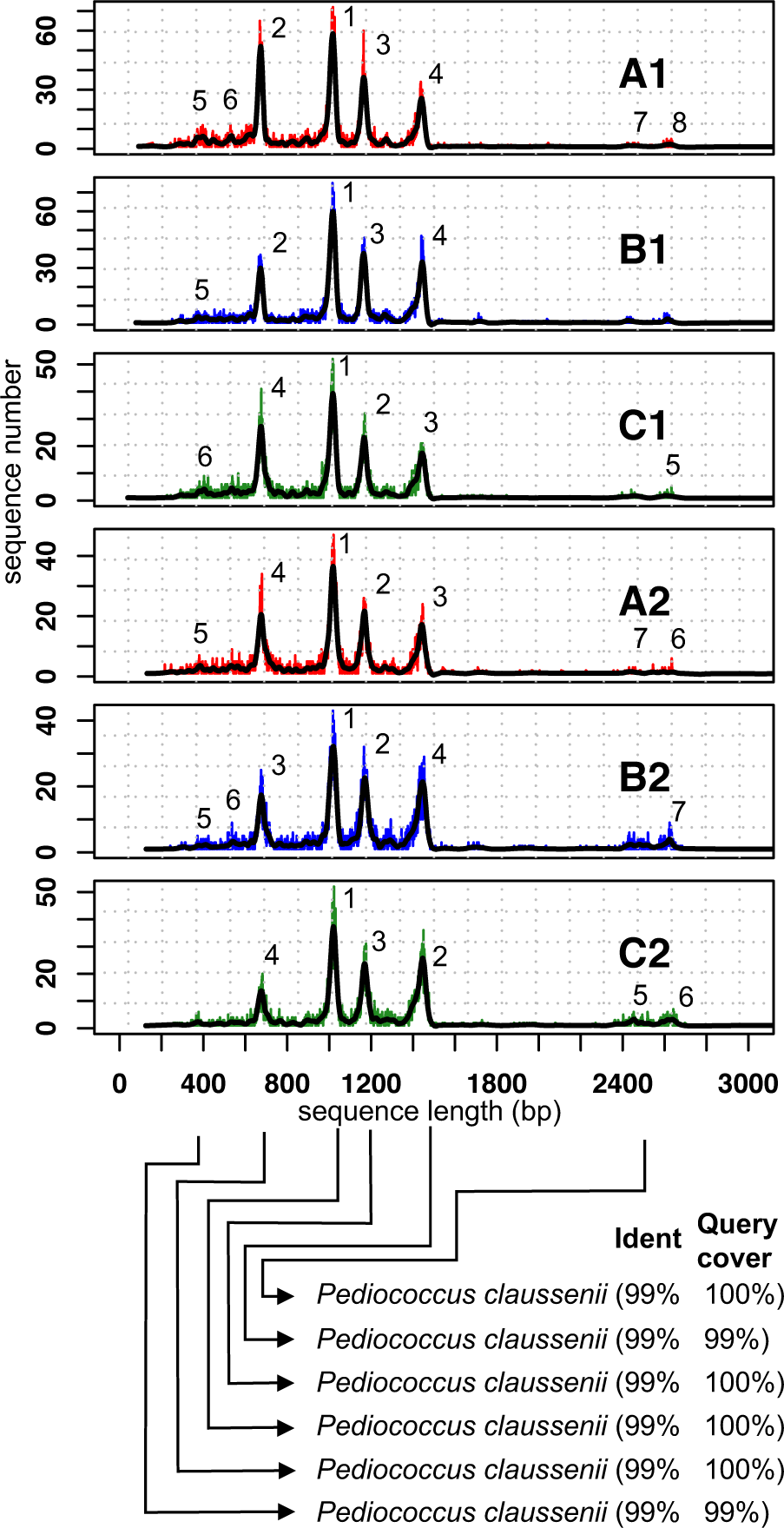
*Pediococcus claussenii* ON-rep-seq LCp. Read length counts profiles (LCp) generated from Oxford Nanopore Technology based rep-PCR amplicon sequencing (ON-rep-seq) of *Pediococcus claussenii.* All technical replicates of *P. clausenni* profiles show high level of similarity across three consecutive sequencing runs (red, blue and green for run A, B and C respectively) and two technical replicates in each run. The retrieved sequences matching the length of a corresponding peak were subjected for correction using Canu and consensus sequences were verified using blastn. For all profiles six to eight high quality reads could be generated, each with > 99% similarity to the reference genome of *P. clausenni.* The numbers above each peak indicates the peak detection sensitivity with 1 being the most evident. The minimum number of reads within the peak needed for reads correction is 50.

### Reads correction within individual peaks provides a set of high quality consensus sequences per isolate allowing detailed identification

A single band on a gel (or peak in LCp) of a Rep-PCR profile will contain mainly representatives of the same amplicon what would allow for base accuracy correction using tools such as e.g. Canu ^22^. With that assumption, we have developed a pipeline operating in three steps: strain specific LCp generation and comparison, within peak reads correction, and peak’s consensus sequence annotation. The pipeline generated on average 10 high quality consensus reads for each isolate (max = 26, min = 3, SD = 4) with mean length of 1 Kbp (max = 3.6 Kbp, min = 0.3 Kbp, SD = 0.6 Kbp). The number of reads used for correction within a peak (cluster size) varied from 50 to 2400 (mean = 254, SD = 246).

Subjecting the set of corrected reads for each sample to centrifuge classifier allowed for unambiguous annotation of all bacteria down to the species and subspecies level (Table 1). The average sequence similarity of corrected reads from strain validated with Illumina sequencing (*S. enterica* serovar Typhimurium C5) reached 99,4% (BLAST; min = 98.3%, max = 100%, SD = 0.5%). Among the isolates tested are for example *Lactobacillus casei* and *Lactobacillus paracasei* subsp. *paracasei* known to be indistinguishable based on 16S rRNA gene sequence comparison or *Lactococcus lactis* subsp. *cremoris* that cannot be distinguished from *Lactococcus lactis* subsp. *lactis*. All these strains were unambiguously discriminated using ON-rep-seq. Two bacterial species: *Bacteroides thetaiotaomicron* and *Lactococcus lactis* subsp. *cremoris* were tested in pairs from different culture collections resulting in all cases in highly reproducible LCp (Supplementary Figure 1).

**Table 01.**
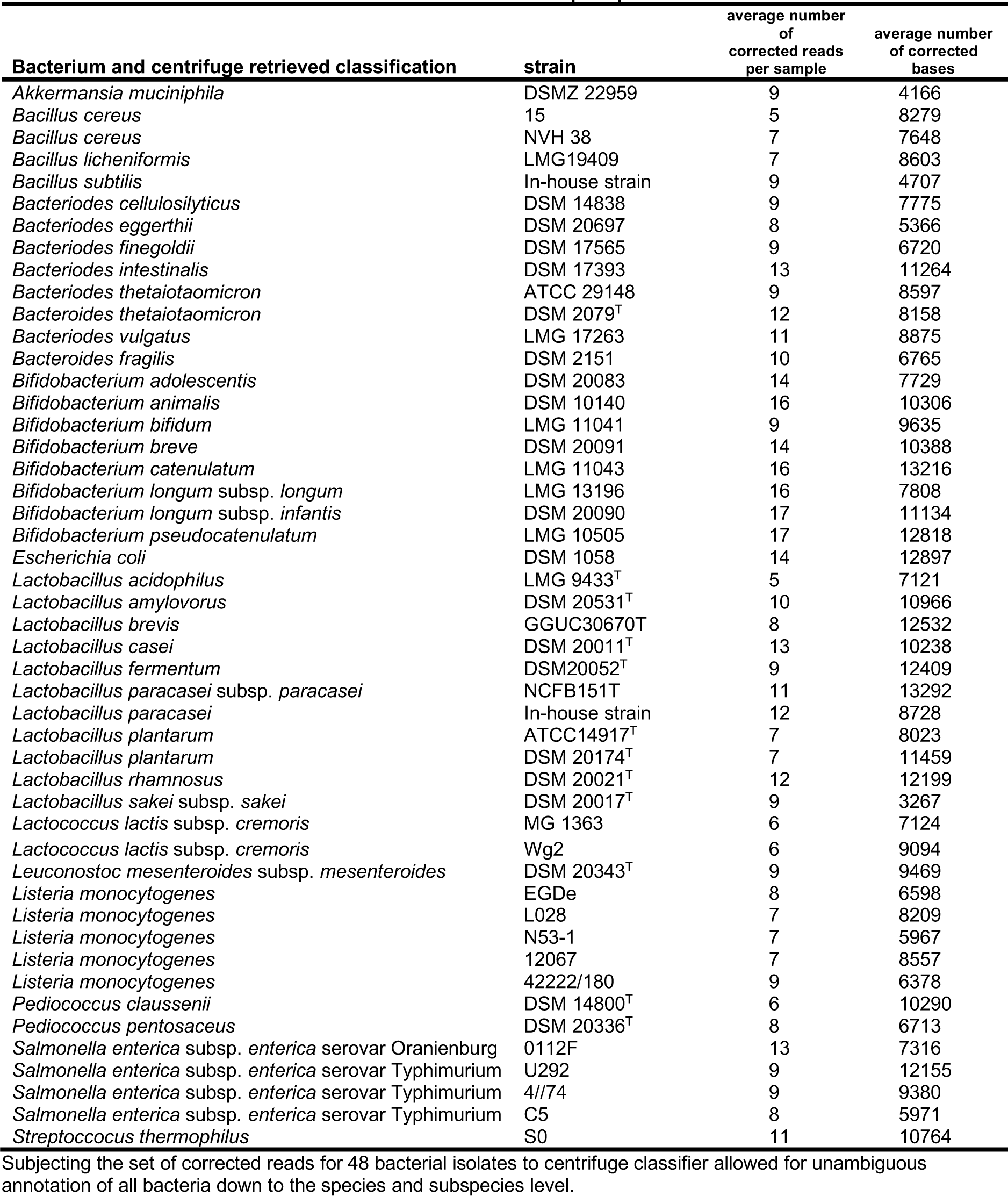
Results of bacterial isolates identification with ON-rep-seq

### Paired comparison of LCp can be used for the strain level differentiation

Five *Listeria monocytogenes*, four *Salmonella enterica* (three serovar Typhimurium and one serovar Oranienburg) and two *Bacillus cereus* strains have been used to evaluate the method for strain level discrimination. We have developed an algorithm (LCpCluster.R) estimating the level of similarity between the pairs of LCp generated by the ON-rep-seq. Among five *L. monocytogenes* strains four unique profiles were identified (Figures 3 and 4). Strains EGDe and LO28 generated identical profiles (Figures 3 A and 4).

**Figure 3.**
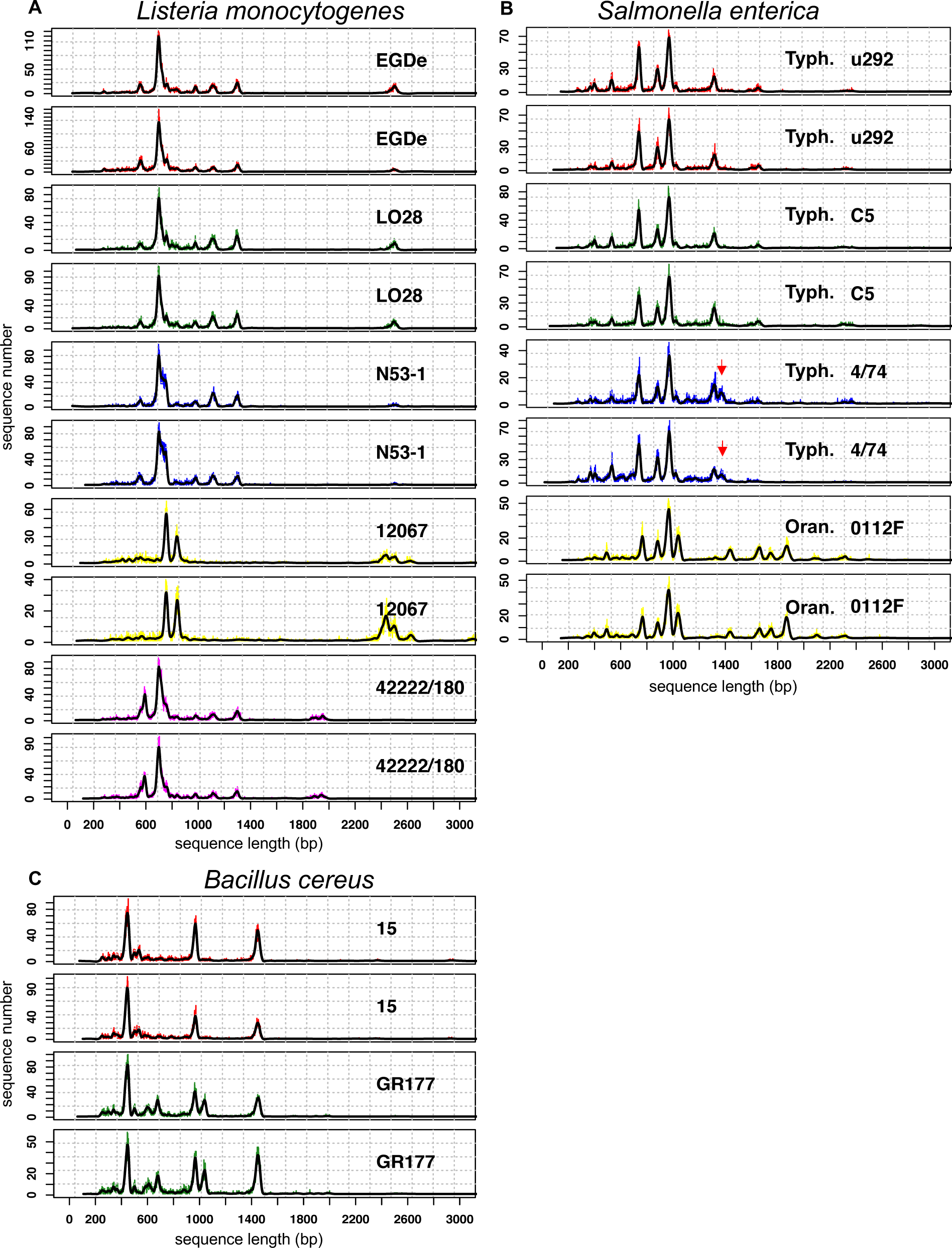
Examples of strain level differentiation using LCp comparison. Oxford Nanopore Technology based rep-PCR amplicon sequencing (ON-rep-seq) of five *L. monocytogenes* (A), four *S. enterica* (B) and two *B. cereus* (C) strains was used to generate read length counts profiles (LCp). All bacterial LCp were produced in duplicates. Consensus sequences from corrected peaks of all 22 samples allowed for unequivocal species and subsbecies level identification. Comparison of LCp revealed four different profiles among the *L. monocytogenes* species. Strains EGDe and LO28 gave highly similar profiles indicating high level of genetical relationship between these two strains (B), what was confirmed by Illumina based shotgun sequencing (orthoANI = 99.9%). Similarly C5 and u292 strains of *S. typhimurium* showed the same profiles (orthoANI = 99.9%) while two other strains could be classified as different (B). The red arrows indicate additional peak distinguishing the 4/74 strain from u292 and C5 that was shown to have a prophage origin. The presence of additional peaks in the LCp of GR177 strain allowed for unambiguous differentiation between the two *B. cereus* strains (C).

**Figure 4.**
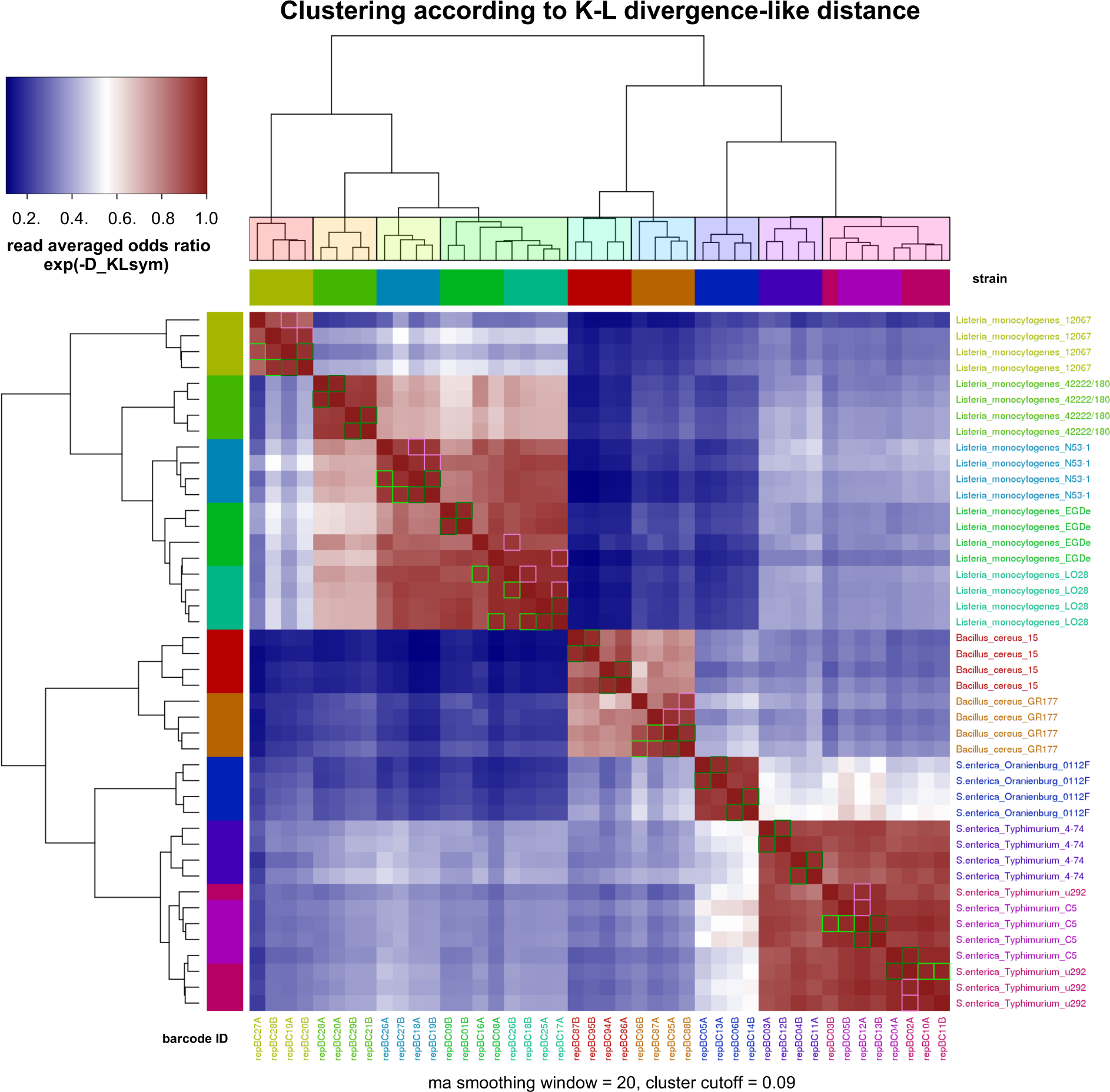
Row/Column clustering according to “Ward.D2” hierarchical clustering on D_KLsym distance. Heatmap showing similarity (10^(-D_KLsym)), and clusters according to cutoff=0.09. Analysis of five Listeria monocytogenes, two Bacillus cereus and four Salmonella enterica serovar Typhimurium strains allowed for species level differentiation in all cases and for strain level differentiation in 8 out of 11 cases. Notably, the presence of the additional peak allowed for unambiguous differentiation of 4/74 from C5 and u292 what was not possible using OrthoANI and MLST analysis based on WGS data. Strain labels colors according to accepted strain similarity derived from visual inspection of profiles in agreement with clustering colors at selected cutoff.

No SNPs variants could be detected when comparing consensus sequences of corresponding peaks of all technical replicates (data not shown). Whole genome sequencing (WGS) data have been used to estimate the genetic similarity between EGDe and LO28 strains. The average nucleotide identity (OrthoANI) index between these two genomes reached 99.9%, while *L. monocytogenes* MLST schemes mapped against EGDe and LO28 found only one differing locus (dapE) out of seven tested (Supplementary Table 1).

This implies a high level of genetical similarity between the two strains that requires a specific approach to ensure differentiation.

Among four *S. enterica* strains LCpCluster.R recognized three unique profiles (Figure 3 B and 4). Serovar Typhimurium strains u292 and C5 showed the same ON-rep-seq LCp with no SNPs variants in corresponding peaks (data not shown). WGS comparison of these two strains revealed high OrthoANI reaching 99.9%. *Salmonella enterica* MLST schemes mapped against genomes of u292 and C5 also showed the same alleles profiles (Supplementary Table 1). This implies that *S. enterica* strains u292 and C5 could not be straightforwardly distinguished based on their genome using both methods.

Interestingly, serovar Typhimurium strain 4/74 that presented similar LCp to u292 and C5, yet with a clear additional peak in the position ∼1370bp (Figure 3 B and 4) reached OrthoANI above 99.9% and had the same MLST profile compared to u292 and C5. In this particular example ON-rep-seq presented higher discrimination power over OrthoANI and MLST analysis based on WGS data. Further investigation of the peak at position ∼1370bp disclosed that the consensus sequence presented high similarity (blast identity 1372/1384bp; 99.1%) to SopEΦ prophage. Moreover, this sequence could only be found in the draft genome of the 4/74 strain (blast identity 1371/1384bp; 99.1%) but not in any of the remaining *S. enterica* strains.

Finally, the two *B. cereus* strains generated clearly distinctive LCp and were classified as different strains. (Figure 3 C and 4). The LCp. Cluster results showing grouping according to “Ward.D2” hierarchical clustering on D_KLsym distance of all 48 isolates in four technical replicates from first two runs are given in Supplementary Figure 2.

### Benchmarking of the R9.4.1 flow cell indicates the theoretical throughput of ON-rep-seq over 1000 bacterial isolates

To validate the method, two R9.4.1 flow cells were benchmarked for the maximum possible output generated. The first benchmarked flow cell generated in total over 2.6 M reads (after quality control and demultiplexing). See Supplementary Table 2 for details. In the first four consecutive runs, each lasting 4h, intertwined with the flow cell washing steps and storage for minimum 24h enough data was generated to successfully demultiplex and identify 4×96 bacterial profiles on a single flow cell. The last run generated 0.22 M reads which was enough to detect and correct sequences of 94 out of 96 samples.

The second flow cell generated in total 2.49 M reads respectively 1 M 0.56 M and 0.87 M for first (4h) second (4h) and third (12h) run. See Table Supplementary Table 2. All three runs of the second flow cell generated enough data to successfully analyze 96 bacterial profiles. To verify the minimum number of reads necessary to analyze all samples the data have been iteratively subsampled and subjected to the analysis with a receiver operating characteristic (ROC) curves to quantify tradeoff between pairwise “same/not-the-same” strain discrimination dependent on clustering cutoff. Throughout the analysis it was noticed that within-strain variance was larger than between-strain variance in cases of small differing features in the latter, and the disproportion of short reads vs long reads in the former case (the observation verified by sample mean read length regression vs sample read count; Supplementary Figure 3). This this disproportion was attributed to the third sequencing run on the reused flow cell, hence the latest repC run was omitted from the cluster analysis and most of ROC curves analysis that follows.

Clustering on different data sets were compared: all, “wo.rep*C” (without the third consecutive run: repC), 2%, 10%, 20%, 50% subsamples. “wo.rep*C” performed best most of the time, though random fluctuations in 50%, 20% and 10% subsamples overperformed occasionally at single data points. Subsampling to 50% and 20% (avg. #reads/sample 4326 and 1730) performed very similarly to full samples (avg. #reads/s 8652), while 10% subsamples performed worse, though still reasonably good, while 2% subsamples (avg. #reads/s 173) performed much worse, though relevant information is still present and retrievable even with such a small reads lengths sample (Supplementary Figure 4 C-H).

The flow cell benchmarking results showed that 20% of generated reads (avg. #reads/sample 1730) were already sufficient to analyze all samples. Notably, the number of isolates that could be analyzed simultaneously on a single flow cell will ultimately depend on the number and position of peaks in LCp (for strain level comparison). Nonetheless, our data demonstrate that the theoretical throughput of the R9.4.1 flow cell ranges between 960 and 1440 isolates depending on the sequencing run performance (∼1.5 M to ∼2.5 M reads respectively).

## Discussion

The process of fast and accurate bacterial identification, subtyping and strain level differentiation is of high importance in epidemiology, to recognize infection outbreaks, determine its source or follow highly virulent nosocomial pathogens. It is also desired in the food industry to validate quality and safety and to investigate microbiologically complex communities like many fermented foods. For the past three decades the most commonly used and standardized methods became molecular techniques based on DNA analysis ^14^. Since first described in 1994 Rep-PCR targeting REP and/or repetitive intergenic consensus (ERIC) regions became a widely used methods of DNA typing ^12,17^. Its discriminatory power has been shown multiple times to be superior to many other typing methods including ribotyping ^12,23^, multilocus enzyme electrophoresis ^24,25^, but also biochemical characterization ^26^. Rep-PCR was often shown to have similar or slightly lower discriminatory power than pulse-field gel electrophoresis (PFGE) but was always considered a less laborious and cheaper solution ^27,28,29^. Among several Rep-PCR options (GTG)5-PCR have reported to be the most robust ^30^. Despite well-documented strain level discrimination power, the main pitfall of Rep-PCR is without a doubt its inability for taxonomic identification, without additional analysis such as 16S rRNA gene sequencing, which requires extra laboratory work and significantly increases time and cost of the analysis. Moreover, such strategy relays entirely in the discriminatory power of 16S rRNA gene that does not always allow for species level identification.

The massive leap in DNA sequencing methods made within the last decade heralded the inevitable decline of many “old-fashioned” DNA finger-print based typing methods. A single HiSeq X instrument (IIlumina) has a capacity to sequence about 35,000 average size bacterial genomes with 100 times coverage in a single run (Illumina.com). Yet, notwithstanding the immense potential, this technology is still not meant for fast, routine and cost-effective typing of bacteria. It is mainly due to the high equipment cost, low flexibility requiring collection of multiple samples (from dozens to thousands depending on the platform), relatively long runtime and complex data analysis. The portable, USB powered MinION offered by Oxford Nanopore Technologies is so far the cheapest (∼1000$) sequencing platform on the market. Its main advantage besides the price is the possibility to generate ultra-long reads with the longest ones crossing 1Mb. Nonetheless, there are two main reasons why ONT have not yet become the first choice of a sequencing strategy in many laboratories. First is the relatively high base calling error rate of a single DNA molecule and second, a relatively low throughput compared to many other platforms ^10^. Our method have largely solved these two hindrances, allowing ONT to be exploited for accurate, large scale and detailed identification of bacteria. Highly reproducible amplification of regions flanked by REP elements not only solved the main problem with sequencing redundancy one needs to deal during WGS, but also enabled single base error correction owing it to its random nature. ON-rep-seq offers the well documented discrimination power of the DNA fingerprint analysis, but also for the first time full access to the “hidden information” within each band DNA sequence in quality crossing 99% accuracy. Since each isolate composes on average of 10 corrected consensus reads with an average length of 1Kb this information can be used for highly accurate taxonomic identification. Even if one of the reads would not find a hit in a database, there are still several others to ensure classification. As shown here, all 48 isolates have been accurately assigned to the species and sub-species level. Also, for the first time users will be able to easily determine contaminations in case one or several peaks would turn out to belong to another organism. Lastly, by reducing the number of samples and increasing the coverage one could achieve even higher accuracy of the consensus sequence what could be used to assess presence of SNPs in profiles of closely related strains. This might be an additional source of previously unknown discrimination repository of Rep-PCR in some unique cases.

We have demonstrated here that ON-rep-seq successfully identified and differentiated between *Listeria monocytogenes* and *Salmonella enterica* serovar Typhimurium strains. Both species are of special epidemiological importance and model organisms for host-pathogen infection ^31^ ^32^. Rep-PCR was previously recommended method for subtyping of *L. monocytogenes* and *S. enterica* with a similar discrimination power of PFGE or RAPD ^29,33^. The only two undistinguished pairs of strains were *L. monocytogenes* EGDe from LO28 and *Salmonella enterica* C5 from u292. Paired comparison of their genomes revealed high level of similarity (OrthoANI > 99.9%) further confirmed with the MLST indicating that genetic diversity between these two strains could be allocated in SNPs. Regrettably, none of these SNPs were found by comparing sequence within the peaks. Although ON-rep-seq cannot discriminate between strains that differs solely with SNPs it can be used for fast and cost-effective screening of multiple isolates to select those of identical profiles that should be subjected for deep sequencing saving resources, money and time.

Interestingly ON-rep-seq was shown to be superior to traditional WGS analysis in distinguishing between *S. enterica* u292 and 4/74. The OrthoANI between these two strains reached > 99.9% with identical MLST profiles. This makes it very challenging to differentiate between u292 and 4/74 at the strain level using WGS ^34,35^. However, comparison of ON-rep-seq based LCp allowed for clear and unambiguous differentiation between the two strains. The peak allowing this distinction was shown to be a mobile element with high similarity to a prophage. It was previously demonstrated that large fractions of genetic variation in in *Salmonella* strains is allocated in variable genomic regions and islands that encompass phage insertions ^36^.

Presented in this work barcodes enable accurate tagmentation of 96 isolates, but our data demonstrate that even about 1000 barcodes could be used on a single R9 flow cell. We have benchmarked two R9.4.1 flow cells to estimate the maximum possible output and cost per isolate. Since the flow cell price ranges between 475$-900$ (depending on the bundle offer) the sole cost of sequencing assuming highest output would range between 0.40 – 0.75 $. It is important to mention that the maximum data output will vary depending on the flow cell viability that may be affected by multiple washing steps. Furthermore, it was demonstrated herein that consecutive usage of the flow cell may be a source of an increased run effect. Therefore, the best performance of ON-rep-seq could be achieved if for example 96×10 barcodes were used in a single run lasting for maximum time (48h). It seems however that the new gadget offered by ONT called Flongle, promised to be released this year, could be the most optimal and user-friendly solution for ON-rep-seq (nanoporetech.com). Flongle is an adapter for MinION with one-quarter throughput of a R9 flow cell but price not crossing 100$. This means that up to 3×96 isolates could be analyzed in a single run for about 0.35$ per sample. Naturally, the user could then choose to sequence less isolates but ensure even better coverage.

In summary, we present here the DNA enrichment and barcoding method called ON-rep-seq (from: Oxford Nanopore based Rep-PCR based sequencing) that in combination with ONT sequencing platforms allows for highly cost effective, bulk screening of bacterial isolates with species and strain level resolution. We believe that ON-rep-seq has a potential to become a modern standard molecular based method with multiple applications in research, industry and medicine. By sharing it to other users we are looking forward for thorough validation of many more bacterial species, optimization of sequencing protocols and pipelines. We hope that conjoined effort of multiple users will also allow for the development of ON-rep-seq consensus reads database facilitating in the even faster and simplified identification.

## Methods

### Wet laboratory Rep-PCR-1

Bacterial genomic DNA was extracted using GenElute^™^ Bacterial Genomic Kit (Sigma Life Science, Darmstadt, Germany) according to the manufacturer’s instructions. In total 48 isolates represented by 38 different bacterial species were subjected for the analysis in duplicates in each of the three runs giving six technical replicates per isolate. The barcodes order was shifted during preparation of each library to ensure that every technical replicate is tagged with different barcode sequence. Three strains of *Salmonella enterica*, five *Listeria monocytogenes* and two *Bacillus cereus* strains have been used to evaluate the ability of the method for strain level differentiation. The detailed list of bacteria used for the analysis is given in Table1 while ON-rep-seq LCp in Supplementary Figure 1. The Rep-PCR reaction mix contained 5 μl PCRBIO HiFi buffer (5x), 0.25 μl of PCRBIO HiFi Polymerase (PCR Biosystems Ltd, London, United Kingdom), 4 μl of (GTG)5 primers (5 μM), 1 μl of DNA (∼20ng/μl) and nuclease-free water to a total volume of 25 μl. The Rep-PCR thermal conditions were optimized as follows: Denaturation at 95°C for 5 min; 30 cycles of 95°C for 30 s, 45°C for 1 min and 62°C for 4 min; followed by final elongation at 72°C for 5 min. It is important to note that several polymerases have been tested in order to shorten the elongation time without compromising the longest amplicons. With current settings, the PCR takes less than 3h on SureCycler 8800 (Agilent, CA, USA).

### Barcoding by dual-stage Rep-PCR-2

We have designed 96 ONT compatible barcodes (Supplementary Table 3) with 10bp spacer separating ONT motor protein adapter from the barcode sequence and (GTG)5 pairing region. The spacer was added to ensure higher tolerance for the low-quality at the beginning of the sequence entering the pore and thus higher recovery of barcode sequence. At the same time the spacer sequence was designed to prevent creations of stem-loops in relatively long primers during low temperature annealing step. The Rep-PCR reaction mix contained 12 μl of PCRBIO UltraMix (PCR Biosystems Ltd, London, United Kingdom) 2 μl of corresponding repBC primer (10 μM), 1 μl of PCR product from Rep-PCR-1 and nuclease-free water to a total volume of 25 μl. Incorporation of ONT compatible adapters (Supplementary Table 3) was performed using dual-stage PCR where first 3 cycles provide optimal annealing of (GTG)5 regions, while next 10 cycles allow for best hybridization of full adapters in consecutive cycles: Denaturation 5 min; 3 cycles of 95°C for 30 s, 45°C for 1 min and 62°C for 4 min; followed by 10 cycles of 95°C for 30 s, 65°C for 1 min and 72°C for 4 min and final elongation at 72°C for 5 min.

### Library preparation and ONT-sequencing

After Rep-PCR-2 samples were pooled using 10 μl of each sample. Note that samples were not pooled in equimolar concentration due to expected differences in length of amplified regions between the samples. However, it is advisable to verify the DNA concentration with Qubit^®^ dsDNA BR Assay Kit (Life Technologies, CA, USA) for the quality control of the Rep-PCR-2 step. The measurement was performed with Varioskan Flash Multimode Reader (Thermo Fischer Scientific, MA, USA). Fluorescence was measured at 485/530 nm.

The pooled library was cleaned with AMPure XP beads (Beckman Coulter Genomic, CA, USA) in volumes 100:50 μl respectively. The bead pellet was washed with 80% ethanol and re-suspended in 100 μl of nuclease-free water. The bead washing step was added to shift the proportion of short to long reads that is multi-template PCR specific feature and to remove primer-dimers. The pooled and bead-purified library was measured with Qubit^®^ dsDNA HS Assay Kit (Life Technologies, CA, USA) and 66 ng of library was used as an input to the End-prep step in 1D amplicon by ligation protocol (ADE_9003_v108_ revT_18Oct2016) with one adjustment: 80% ethanol instead of 70% was used for all washing steps.

To validate our method, we have benchmarked two R9.4.1 flow cells for the maximum possible output generated. First flow cell was used 10 days after the delivery in five consecutive runs, each lasting 4h, intertwined with the flow cell washing steps and storage for minimum 24h (QC 1347 active pores). Second flow cell was used 44 days after the delivery in three consecutive runs lasting respectively 4 h, 4h and 12h (to collect maximum amount of data from declining flow cell; QC 1105 active pores). After each run the flow cell was washed according to the manufacturer’s instructions and a new library was prepared and loaded. In order to evaluate the possibility of barcode specific amplification during Rep-PCR-2 step, all samples received different barcode in consecutive sequencing runs. The data from the first benchmarked flow cell were used solely to test the optimal concentration of DNA needed and viability of the flow cell while data from the second flow cell are presented herein and can be downloaded from SRA NCBI repository (#SUB4333515).

## Data analysis

### Data collection, base calling, demultiplexing and trimming

Data were collected using MinKnow 1.10.23. The amount of data collected in both R.9.4 flow cells is listed in Supplementary Table 2. Albacore v2.1.3 pipeline was used to base call raw fast5 to fastq. Porechop v0.2.2 was used for adapters trimming and samples demultiplexing. Porechop settings together with the list of custom adapters (adapters.py) compatible with oligos given in Supplementary Table 3 are available at (bitbucket.org/modelscat/on-rep-seq) The script allows for demultiplexing up to 96 barcodes and trimming of: ONT adapters, custom spacers and tandem repeats of (GTG)n.

### Correction and base location of peaks

Peaks are identified in LCp expressed as sequencing length (x-axis) by number of reads (y-axis) by fitting local third order polynomials in a sliding window of size 1/50 of the x-span across the x-axis, followed by calculation of the first- and second order derivatives. The position of a peak is identified at the x-axis where the first derivative is zero and the second derivative is negative. Only peaks with intensity higher than baseline, defined as a moving boxcar (zero order polynomial) in a broad window (4 times the size of the window used for calculation of the derivative) are used for further analysis. The identified peaks are ordered based on the height, and a representative fragment are used for data base matching.

### Reads correction within a peak

Sequences containing quality scores (fastq files) resolved within each peak were retrieved using Cutadapt v1.15 ^37^, and corrected with Canu v1.6 ^22^ using the following parameters: genomeSize=5k, minimumReadLength=200, correctedErrorRate=0.05, corOutCoverage=5000, corMinCoverage=2 and minOverlapLength=50. The corrected reads were sorted-by-length and clustered with UCLUST (cluster_fast) from USEARCH v10.0 ^38^, using the following options: -id of 0.9, -minsl of 0.8, -sizeout, and min_cons_pct of 20. Subsequently, consensus sequences were sorted-by-size (coverage) and those with a minimum coverage-size of 50× were kept for downstream analyses.

### Classification

Centrifuge 1.0.3 microbial classification engine was used for labelling of corrected reads.

### Comparison of LCp

The identification of a good distance measure on read length counts profiles (LCp) was approached by considering them as approximating samples of their underlying sampling distributions. Ideally one would like to understand processes involved in signal peaks and noise formation, thus a priori distributions could be postulated, and later optimized for profiles posteriors. Primarily, empirical discrete length distributions were smoothed with window moving average (ma). Selection of ma window size was done by computing the average jitter of all profiles: an average number of times when profile’s discrete derivative changes sign (change to 0 was counted as 0.5). From mean jitter plot “ma” window size was selected to 20, the point of the lowest second derivative, after which second derivative stabilized closely around 0, meaning the information (jitter) loss due to increasing of window size became relatively low and more constant (Supplementary Figure 4 A, B). Next with each LCp assigned was a “ma” smoothed and probability-normalized “distribution profile” Dp.

Stability problems around #reads(i)=Dp(i)=0 are avoided by considering a mixture of “ma” smoothed Dp and the uniform read lengths distribution (in a considered range 150 to 3000bp) with proportions (0.99, 0.01). The distance between two samples, reads lengths-based, was defined as a function of LCp_1, Dp_1, LCp_2, Dp_2. One natural approach was to consider the probability of sampling LCp_1 from Dp_2, however for the distance to be comparable between samples of different read counts it needed to be normalized by total read count. Resulting is the following logarithmic formula:

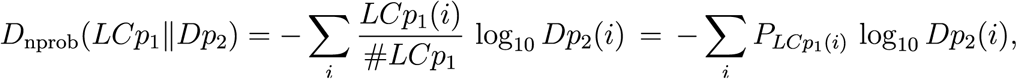

The above formula is however not centralized because the distance of a sample to itself is not 0 but it is rather equal to sample’s smoothed entropy. Centralization of this distance yields distance very similar to Kullback-Leiber divergence of probabilities, which is proposed for the distance between LCp, as follows:

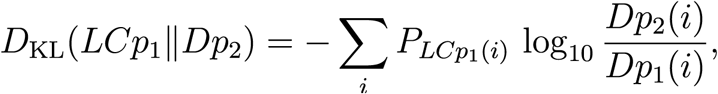

In the following clustering analysis, we use the symmetrized version:

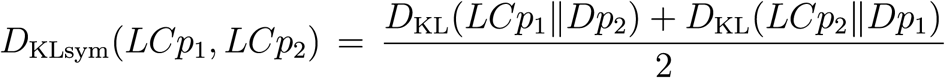

### Analysis of D_KLsym distance between peaks profiles performance on bacterial LCp generated with ON-rep-seq

Validation of KL based distance on LCp by hierarchical clustering was performed on sequencing results where clusters were compared with down-to-strain sample labels. To promote clusters with low variance around centroids “Ward.D2” clustering method was selected and performed with modified “heatmap3” R library. Figure 4 shows clusters recovered with cutoff=0.09 where all *L. monocytogenes, B. cereus* and *S. enterica* strains with clearly visible feature peaks were properly clustered.

### Whole genome sequencing data analysis

Complete or draft genomes *L. monocytogenes* EGDe (#NC_003210.1) and LO28 (#AARY02000001.1-2001127.1); *S. enterica* serovar Typhimurium ST4/74 (#CP002487.1) and u292 (#ERR277220) were downloaded from public databases and compared using OrthoANI ^8^.

*Salmonella enterica* MLST schemes (internal fragments and their alleles) hosted at PubMLST.org were mapped against genomes of U292 (#ERR277220) and 4/74 ((#CP002487.1), as well as the assemble contigs of C5 strain included in this study.

For strain C5, DNA was subjected to library preparation (Nextera XT kit, following manufacturer procedures) and sequencing on Illumina NextSeq platform. High-quality reads (>95% quality and minimum size of 50nt using Trimmomatic v0.35 ^39^ were de-convoluted from phiX174 controls reads (-id: 0.97, -query_cov: 0.97) and dereplicated using USEARCH v10 ^40^. Subsequently, reads were assembled into contigs using Spades v3.5.0 ^41^. Contigs with a minimum size of 10,000 bp generated for C5 strain, and in addition to the publicly available U292 and 4/74 putative genomes, were subjected to MLST analysis on the CLC Genomics Workbench v11.1 using a minimum alignment length of 400bp and high level of alignment stringency.

## Acknowledgements

To **Pernille Johansen** for inspirational conversation. To **Witold Kot** and Aarhus University for generating WGS data. To **Basheer Yousef Aideh** for providing access to the broad bacterial collection. To **Henrik Siegumfeldt** for help in editing.

## Authors contributions

L.K conceived the presented idea, participated in the data analysis, developed the wet-lab protocol, participated in pipeline development, designed figures, wrote the manuscript. J.L.C.M participated in pipeline development, data analysis and interpretation, tables generation, contributed in manuscript writing. D.N.M participated in optimization of the wet-lab protocol, data analysis and interpretation. M.B.M participated in optimization of the wet-lab protocol. M.A.R participated in the data analysis and interpretation, pipeline development, contributed in manuscript writing. M.S participated in the data analysis, interpretation, pipeline development, designed several figures, contributed in manuscript writing. D.S.N supervised the work, participated in formulation of the initial idea, contributed in manuscript writing. All authors edited and approved the final version of the manuscript.

## Corresponding authors

Lukasz Krych or Josué L. Castro-Mejia or Dennis S. Nielsen

## Software and code

ON-rep-seq pipeline is available at: bitbucket.org/modelscat/on-rep-seq

## Data

Fastq files can be downloaded from SRA NCBI repository (#SUB4333515)

---

**Supplementary Figure 1.**
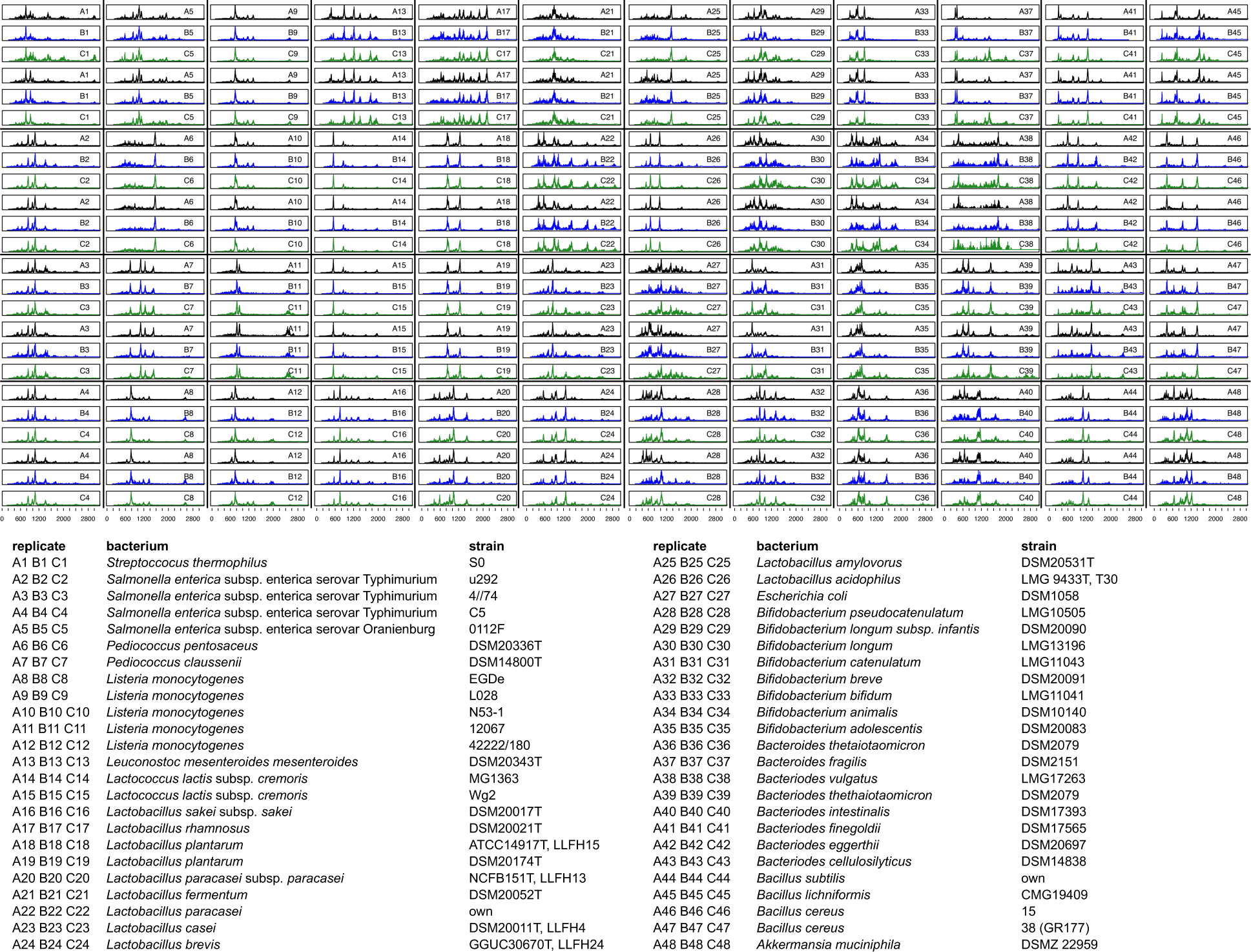
LCp generated for 48 bacterial isolates using Oxford Nanopore Technology based rep-PCR amplicon sequencing (ON-rep-seq). The black, blue and green profiles indicate data collected during run A, B and C respectively for which each technical replicate received different barcode. All isolates were analysed in duplicates within each run. The list of bacterial taxa matching given LCp is given in the table.

**Supplementary Figure 2.**
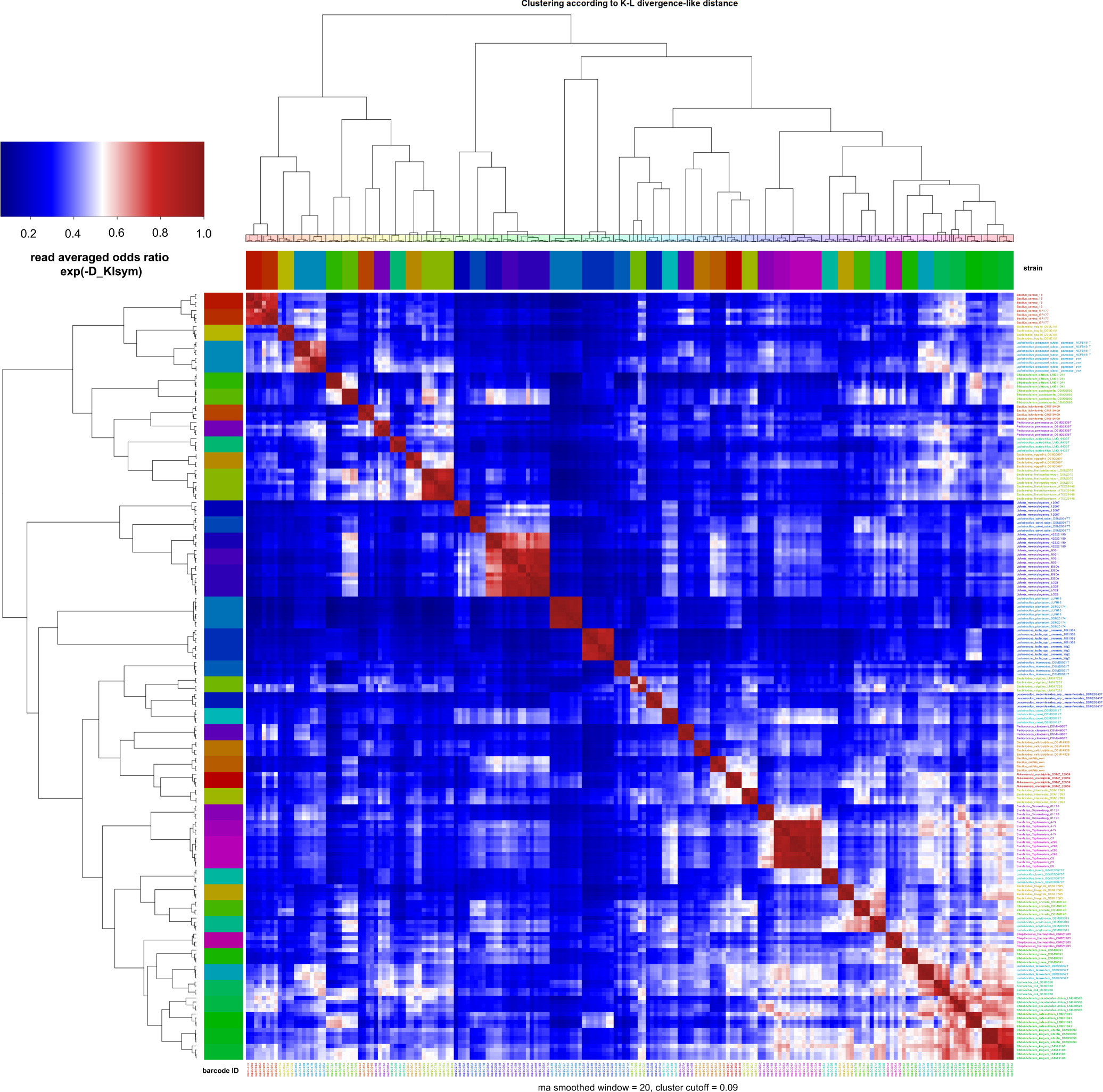
Row/Column clustering according to “Ward.D2” hierarchical clustering on D_KLsym distance of all 48 isolates. Heatmap showing similarity (10^(-D_KLsym)), and clustering according to cutoff=0.09. The detailed analysis using varying cutoff value (no single cutoff achieves exact separation between all and only different LCp, see Figure S4 C, D ROC curves) and LCp visual inspection allowed for accurate differentiation between all except two pairs of bacterial strains described thoroughly in the results section (see Figure 3 and Figure S2 for details). Technical replicates from the third run “repC” were removed from the analysis due to higher short/long reads imbalance.

**Supplementary Figure 3.**
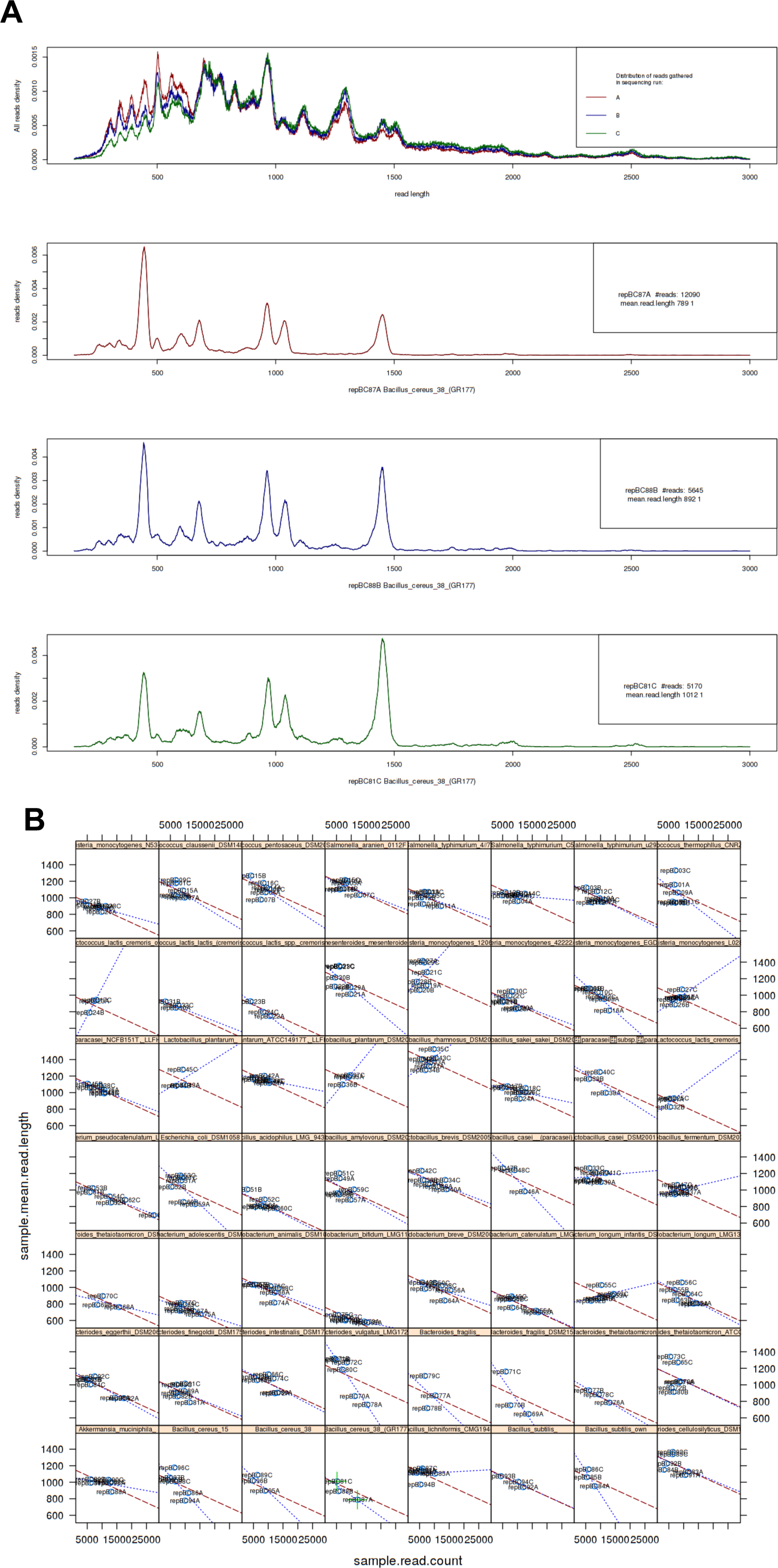
A)Top panel presents distribution of lengths of reads obtained in 3 separate consecutive sequencing runs A, B, C on the same flow cell. Third run C obtained less short reads, some differences are also visible in second run B, compared to the first run A. Bottom 3 panels show LCps of Bacillus_cereus_38(GR177) strain obtained from runs A,B,C. B) Regression analysis of mean read length from LCp vs read count in LCp, data shown in separate panels for each strain replicates. Red dashed line is regression line obtained in all samples analysis, blue lines are regression lines for each strain only. Green markers mark runs A, C for Bacillus_cereus_38(GR177) (panels 2 and 4 in A).

**Supplementary Figure 4.**
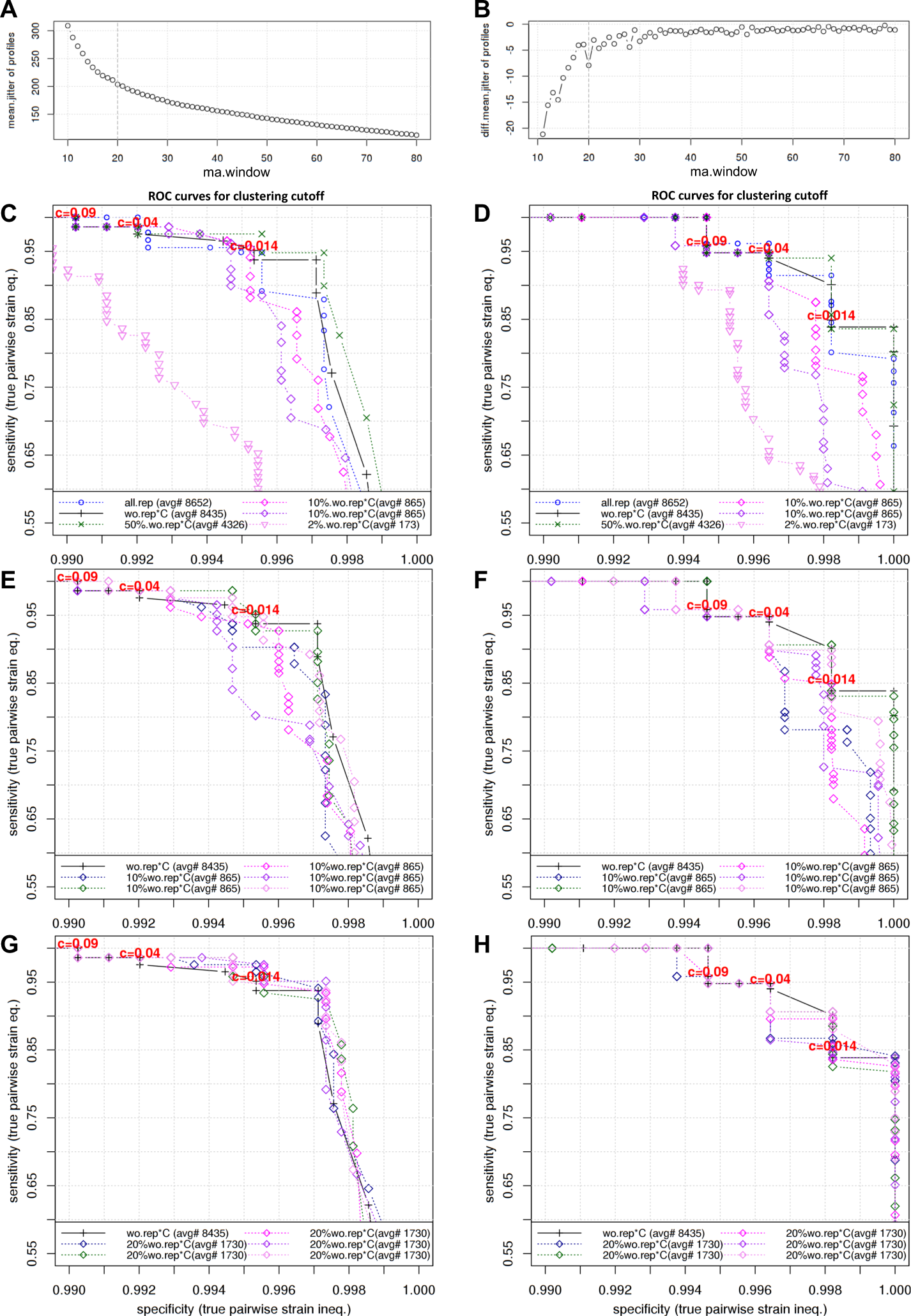
Peaks profiles comparison. A) mean jitter of all profiles dependence on smoothing moving average “ma” window size. Jitter was defined as an average number of times when profile’s discrete derivative changes sign (change to 0 was counted as 0.5). B) discrete derivative (diff lag=1) of the (top) mean jitter. Mean jitter changes more slowly and steadily with sizes of ma.window > 20, suggestive of stabilization (noise decoupling) of information content in larger smoothing window results.C-H) Receiver operating characteristic (ROC) curves of pairwise “same/not-the-same” strain discrimination in various cutoffs c (diff. step=0.005), for various subsets of data: “all”, “wo.rep*C” dataset without the third sequencing run “C” on twice used flow cells, “50%.wo.rep*C” subsample half the size of original, “20%.wo.rep*C” five subsamples 1/20^th^ of reads, “10%.wo.rep*C” seven subsamples 1/10^th^ of reads and “2%.wo.rep*C” 1/50^th^ of reads. On x-axis specificity, the percentage of correctly identified “not-the-same strain” pairs out of all such pairs (36096 for wo.rep*C), on y-axis sensitivity, the percentage of correctly identified “same strain” pairs, out of all such pairs (768 for wo.rep*C). C) Clustering according to sample strain label, which can be thought of more as a whole method performance, in contrast to D. Some values on the plot: c=0.09 (sp 0.9902, se 1.0), c=0.014 (sp 0.9953, se 0.9514). D) Clustering according to sample strain similarity derived from visual inspection of profiles, thus these curves correspond more to D_KLsym-based profile comparison performance, than to the whole method. Values on the plot: c=0.09 (sp 0.9947, se 0.9583), c=0.014 (sp 0.9982, se 0.8490). All cutoffs “c” values marked for “wo.rep*C”. E) Clustering according to sample strain label using 5 iterations of 10% subsets. F) Clustering according to sample strain similarity derived from visual inspection of profiles using 5 iterations of 20% subsets. The analysis shows that 20% subsets perform similarly to the whole dataset what indicates the theoretical throughput of ON-rep-seq to range from 960 (if generating ∼1.5M reads) to 1440 (if generating ∼2.5M reads) isolates per flow cell.

**Supplementary Table 1.** Identification of MLST genes alleles among selected strains of *Salmonella enterica* subsp. *enterica* serovar Typhimurium and *Listeria monocytogenes*

| Strain | Gene | Allele type | Strain | Gene | Allele type |
| --- | --- | --- | --- | --- | --- |
| *Salmonella enterica*<br>U292 | aroC | 10 | *Listeria monocytogenes*<br>EDGe | abcZ | 6 |
|  | dnaN | 7 |  | bglA | 5 |
|  | hemD | 12 |  | cat | 6 |
|  | hisD | 9 |  | **dapE** | **20** |
|  | purE | 5 |  | dat | 176 |
|  | sucA | 9 |  | ldh | 4 |
|  | thrA | 2 |  | lhk | 1 |
| *Salmonella enterica*<br>C5 | aroC | 10 | *Listeria monocytogenes*<br>L028 | abcZ | 6 |
|  | dnaN | 7 |  | bglA | 5 |
|  | hemD | 12 |  | cat | 6 |
|  | hisD | 9 |  | **dapE** | **51** |
|  | purE | 5 |  | dat | 176 |
|  | sucA | 9 |  | ldh | 4 |
|  | thrA | 2 |  | lhk | 1 |
| *Salmonella enterica*<br>4/74 | aroC | 10 |  |  |  |
|  | dnaN | 7 |  |  |  |
|  | hemD | 12 |  |  |  |
|  | hisD | 9 |  |  |  |
|  | purE | 5 |  |  |  |
|  | sucA | 9 |  |  |  |
|  | thrA | 2 |  |  |  |

**Supplementary Table 2.** Details regarding benchmarking of two R9.4.1 flow cells.

| Run ID | Flow cell 1 |  |  |  | Flow cell 2 |  |  |
| --- | --- | --- | --- | --- | --- | --- | --- |
|  | A | B | C | D | A | B | C |
| Run Time (h) | 4 | 4 | 4 | 4 | 4 | 4 | 12 |
| Break between the next run (day) | 1 | 4 | 3 | 7 | 1 | 1 | 1 |
| Active pores at start | 1347 | 1324 | 1098 | 925 | 1034 | 779 | 615 |
| Voltage at start (mV) | -180 | -180 | -190 | -195 | -180 | -180 | -190 |
| Initial sequences in strand | ~300 | ~200 | ~150 | ~50 | ~200 | ~120 | ~70 |
| Total number of high quality reads collected | $9.4 \cdot 10^5$ | $7.9 \cdot 10^5$ | $5.7 \cdot 10^5$ | $2.2 \cdot 10^5$ | $10.5 \cdot 10^5$ | $5.7 \cdot 10^5$ | $8.7 \cdot 10^5$ |
| Library concentration loaded in 12 μl (ng/ μl) | 2.5 | 1.8 | 3.0 | 1.6 | 3.2 | 2.1 | 2.4 |

Both flow cells generated in total similar amount of data, although flow cell 1 was in much better condition and had more active pores at start what allowed to perform four consecutive runs. Flow cell 2 had lower number of active pores at arrival and seemed to deteriorate faster therefore only three runs were conducted. Last run was elongated to 12 h in order to collect maximum amount of data from declining flow cell. The data from the first benchmarked flow cell were used solely to test the optimal concentration of DNA needed and viability of the flow cell while data from the second flow cell are presented herein

**Supplementary Table 3.**
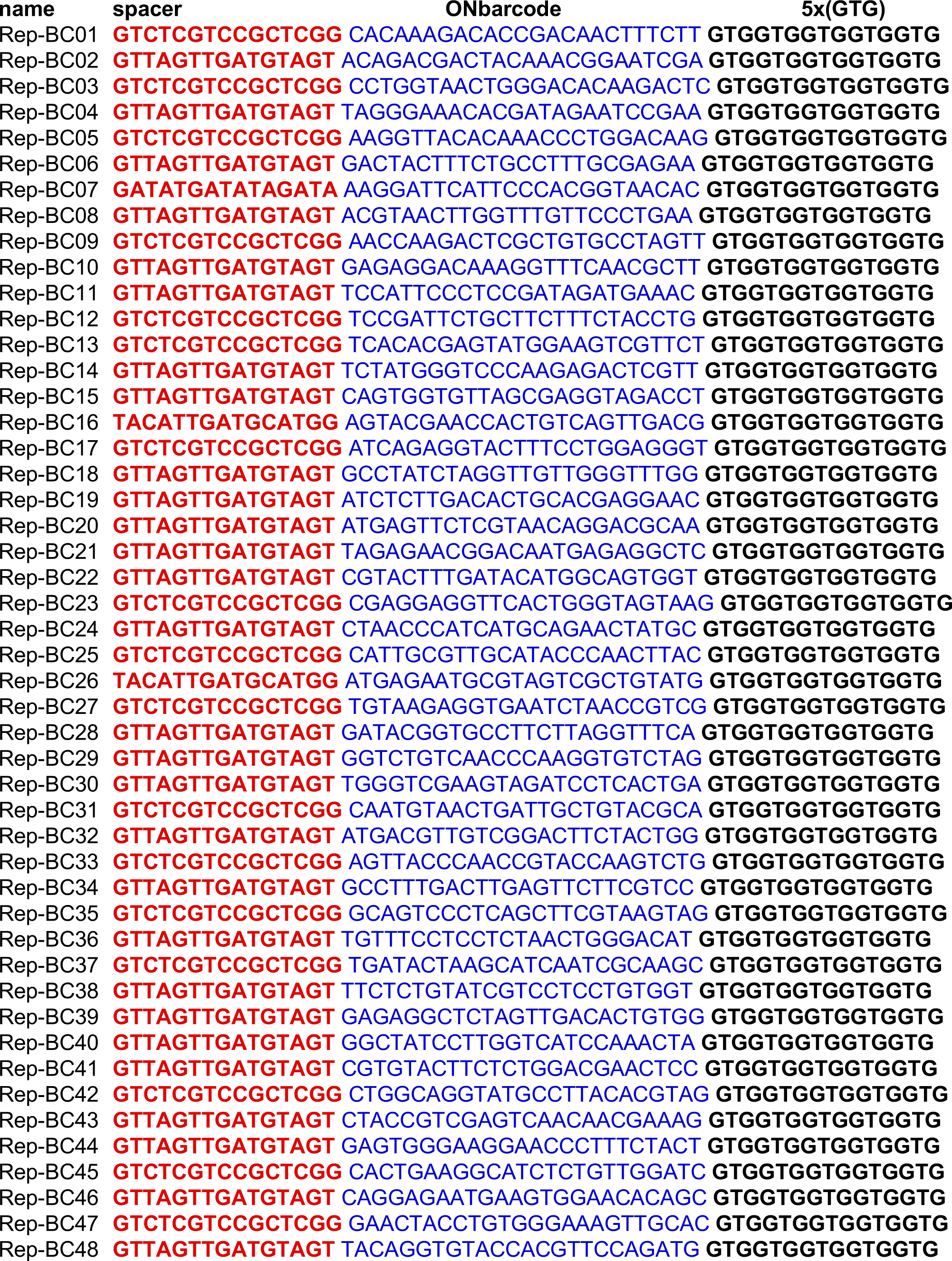
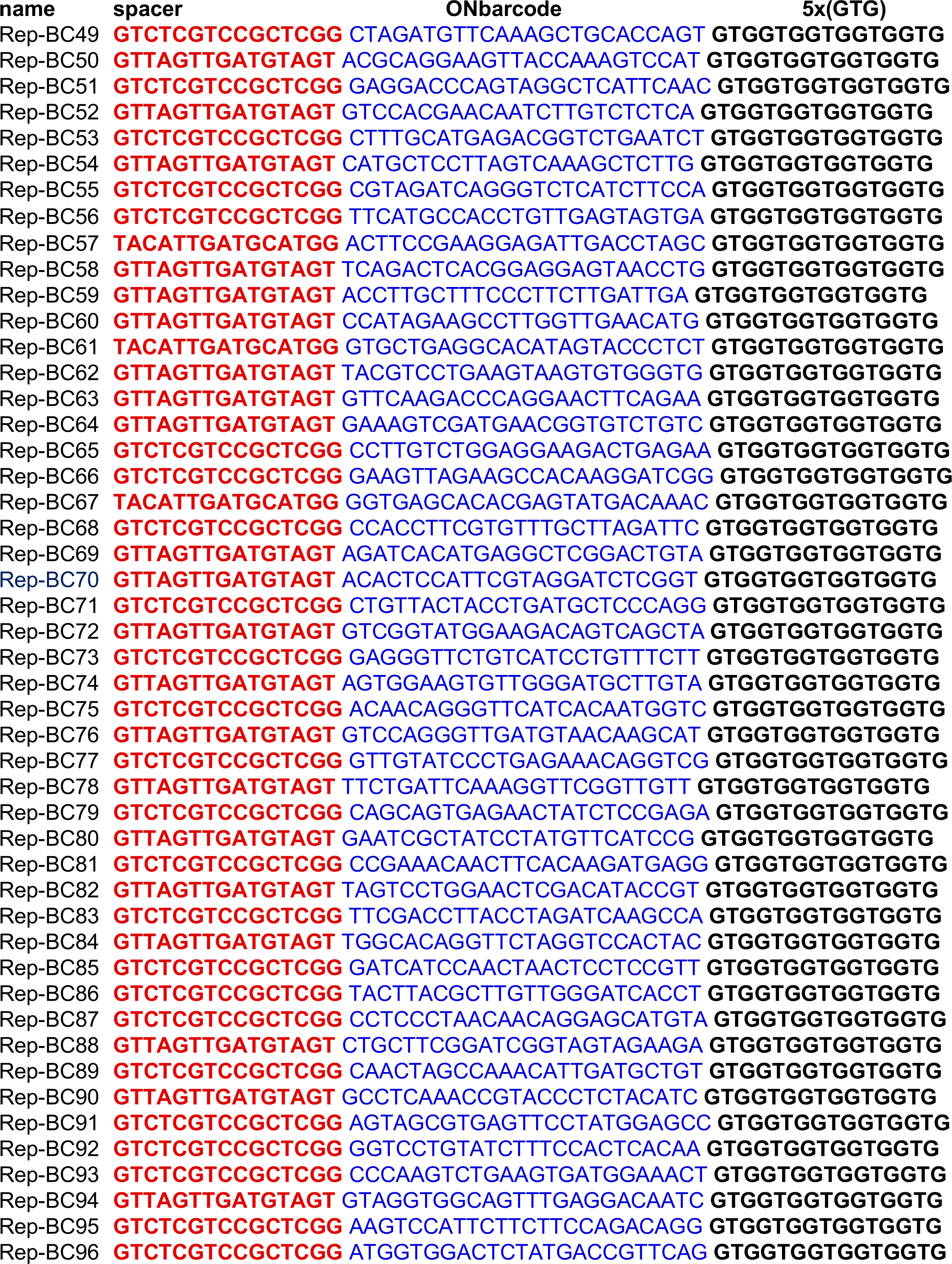
List of 96 barcodes for bacterial isolate Rep-PCR amplicon tagmentation.

## References

1. Marx, V. Microbiology: The road to strain-level identification. Nat. Methods 13, 401–404 (2016).

2. Janda, J. M. & Abbott, S. L. 16S rRNA gene sequencing for bacterial identification in the diagnostic laboratory: Pluses, perils, and pitfalls. Journal of Clinical Microbiology 45, 2761–2764 (2007).

3. Hrabák, J., Chudáčková, E. & Walková, R. Matrix-assisted laser desorption ionization-time of flight (MALDITOF) mass spectrometry for detection of antibiotic resistance mechanisms: From research to routine diagnosis. Clin. Microbiol. Rev. 26, 103–114 (2013).

4. Rodriguez, M. et al. Discriminatory Indices of Typing Methods for Epidemiologic Analysis of Contemporary Staphylococcus aureus Strains. Med. 94, e1534 (2015).

5. Sandrin, T. R., Goldstein, J. E. & Schumaker, S. MALDI TOF MS profiling of bacteria at the strain level: A review. Mass Spectrometry Reviews 32, 188–217 (2013).

6. Miller, J. R. et al. Hybrid assembly with long and short reads improves discovery of gene family expansions. BMC Genomics 18, (2017).

7. Carlisle, E. M. et al. Murine Gut Microbiota and Transcriptome Are Diet Dependent. Ann. Surg. 257, 1 (2012).

8. Lee, I., Kim, Y. O., Park, S. C. & Chun, J. OrthoANI: An improved algorithm and software for calculating average nucleotide identity. Int. J. Syst. Evol. Microbiol. 66, 1100–1103 (2016).

9. Scholz, M. et al. Strain-level microbial epidemiology and population genomics from shotgun metagenomics. Nat. Methods 13, 435–438 (2016).

10. Laver, T. et al. Assessing the performance of the Oxford Nanopore Technologies MinION. Biomol. Detect. Quantif. 3, 1–8 (2015).

11. Stern, M. J., Ames, G. F. L., Smith, N. H., Clare Robinson, E. & Higgins, C. F. Repetitive extragenic palindromic sequences: A major component of the bacterial genome. Cell 37, 1015–1026 (1984).

12. Versalovic, J., Schneider, M. & Bruijn, F. De. Genomic fingerprinting of bacteria using repetitive sequence-based polymerase chain reaction. Methods Mol. Cell. Biol. 25–40 (1994).

13. Versalovic, J., Woods, C. R., Georghiou, P. R., Hamill, R. J. & Lupski, J. R. DNA-based identification and epidemiologic typing of bacterial pathogens. Archives of Pathology and Laboratory Medicine 117, 1088–1098 (1993).

14. Olive, D. M. & Bean, P. Principles and applications of ligation mediated PCR methods for DNA-based typing of microbial organisms. J. Clin. Microbiol. 37, 1661–1669 (1999).

15. De Vuyst, L. et al. Validation of the (GTG)5-rep-PCR fingerprinting technique for rapid classification and identification of acetic acid bacteria, with a focus on isolates from Ghanaian fermented cocoa beans. Int. J. Food Microbiol. 125, 79–90 (2008).

16. Gevers, D., Huys, G. & Swings, J. Applicability of rep-PCR fingerprinting for identification of Lactobacillus species. FEMS Microbiol. Lett. 205, 31–36 (2001).

17. Ishii, S. & Sadowsky, M. J. Applications of the rep-PCR DNA fingerprinting technique to study microbial diversity, ecology and evolution. Environ. Microbiol. 11, 733–740 (2009).

18. Tafvizi, F. & Tajabadi Ebrahimi, M. Application of repetitive extragenic palindromic elements based on PCR in detection of genetic relationship of lactic acid bacteria species isolated from traditional fermented food products. J. Agric. Sci. Technol. 17, 87–98 (2015).

19. Wise, M. G. et al. Predicting Salmonella enterica serotypes by repetitive sequence-based PCR. J. Microbiol. Methods 76, 18–24 (2009).

20. Nurhayati, Priyambada I. D., Radjasa, O. K. & Widada, J. Repetitive element palindromic PCR (Rep-PCR) as a genetic tool to study diversity in amylolytic bacteria. Adv. Sci. Lett. 23, 6458–6461 (2017).

21. Healy, M. et al. Microbial DNA typing by automated repetitive-sequence-based PCR. J. Clin. Microbiol. 43, 199–207 (2005).

22. Koren, S. et al. Canu: Scalable and accurate long-read assembly via adaptive κ-mer weighting and repeat separation. Genome Res. 27, 722–736 (2017).

23. Appuhamy, S., Parton, R., Coote, J. G. & Gibbs, H. A. Genomic fingerprinting of Haemophilus somnus by a combination of PCR methods. J. Clin. Microbiol. 35, 288–291 (1997).

24. Woods, C. R., Versalovic, J., Koeuth, T. & Lupski, J. R. Analysis of relationships among isolates of Citrobacter diversus by using DNA fingerprints generated by repetitive sequence-based primers in the polymerase chain reaction. J. Clin. Microbiol. 30, 2921–2929 (1992).

25. Harvey, J., Norwood, D. E. & Gilmour, A. Comparison of repetitive element sequence-based PCR with multilocus enzyme electrophoresis and pulsed field gel electrophoresis for typing Listeria monocytogenes food isolates. Food Microbiol. 21, 305–312 (2004).

26. Clarridge, J. E. et al. Strategy to detect and identify Bartonella species in routine clinical laboratory yields Bartonella henselae from human immunodeficiency virus-positive patient and unique Bartonella strain from His cat. J. Clin. Microbiol. 33, 2107–2113 (1995).

27. Gunawardana, G. A., Townsend, K. M. & Frost, A. J. Molecular characterisation of avian Pasteurella multocida isolates from Australia and Vietnam by REP-PCR and PFGE. Vet. Microbiol. 72, 97–109 (2000).

28. Northey, G., Gal, M., Rahmati, A. & Brazier, J. S. Subtyping of Clostridium difficile PCR ribotype 001 by REP-PCR and PFGE. J. Med. Microbiol. 54, 543–547 (2005).

29. Chou, C. H. & Wang, C. Genetic relatedness between Listeria monocytogenes isolates from seafood and humans using PFGE and REP-PCR. Int. J. Food Microbiol. 110, 135–148 (2006).

30. Mohapatra, B. R. & Mazumder, A. Comparative efficacy of five different rep-PCR methods to discriminate Escherichia coli populations in aquatic environments. Water Sci. Technol. 58, 537–547 (2008).

31. Hamon, M., Bierne, H. & Cossart, P. Listeria monocytogenes: A multifaceted model. Nature Reviews Microbiology 4, 423–434 (2006).

32. Leekitcharoenphon, P., Nielsen, E. M., Kaas, R. S., Lund, O. & Aarestrup, F. M. Evaluation of whole genome sequencing for outbreak detection of salmonella enterica. PLoS One 9, (2014).

33. Albufera, U., Bhugaloo-Vial, P., Issack, M. I. & Jaufeerally-Fakim, Y. Molecular characterization of Salmonella isolates by REP-PCR and RAPD analysis. Infect. Genet. Evol. 9, 322–327 (2009).

34. Konstantinidis, K. T. & Tiedje, J. M. Genomic insights that advance the species definition for prokaryotes. Proc. Natl. Acad. Sci. 102, 2567–2572 (2005).

35. Konstantinidis, K. T., Ramette, A. & Tiedje, J. M. Toward a more robust assessment of intraspecies diversity, using fewer genetic markers. Appl. Environ. Microbiol. 72, 7286–7293 (2006).

36. Jacobsen, A., Hendriksen, R. S., Aaresturp, F. M., Ussery, D. W. & Friis, C. The Salmonella enterica Pan-genome. Microbial Ecology 62, 487–504 (2011).

37. Martin, M. Cutadapt removes adapter sequences from high-throughput sequencing reads. EMBnet.journal 17, 10 (2011).

38. Edgar, R. C. Search and clustering orders of magnitude faster than BLAST. Bioinformatics 26, 2460–2461 (2010).

39. Bolger, A. M., Lohse, M. & Usadel, B. Trimmomatic: A flexible trimmer for Illumina sequence data. Bioinformatics 30, 2114–2120 (2014).

40. Edgar, R. C. Sequence analysis Search and clustering orders of magnitude faster than BLAST. 26, 2460–2461 (2010).

41. Bankevich, A. et al. SPAdes: A New Genome Assembly Algorithm and Its Applications to Single-Cell Sequencing. J. Comput. Biol. 19, 455–477 (2012).

